# Stochastic analysis of frequency bandwidth and noise attenuation in neurotransmission

**DOI:** 10.1101/2022.04.01.486751

**Authors:** Zahra Vahdat, Abhyudai Singh

## Abstract

Action potential (AP)-triggered neurotransmitter release forms the key basis of inter-neuronal communication. We present a stochastic hybrid system model that captures the release of neurotransmitter-filled vesicles from a presynaptic neuron. More specifically, vesicles arrive as a Poisson process to attach at a given number of docking sites, and each docked vesicle has a certain probability of release when an AP is generated in the presynaptic neuron. The released neurotransmitters enhance the membrane potential of the postsynaptic neuron, and this increase is coupled to the continuous exponential decay of the membrane potential. The buildup of potential to a critical threshold level results in an AP firing in the postsynaptic neuron, with the potential subsequently resetting back to its resting level. Our model analysis develops formulas that quantify the fluctuations in the number of released vesicles and mechanistically connects them to fluctuations in both the postsynaptic membrane potential and the AP firing times. Increasing the frequency of APs in the presynaptic neuron leads to saturation effects on the postsynaptic side, resulting in a limiting frequency range of neurotransmission. Interestingly, AP firing in the postsynaptic neuron becomes more precise with increasing AP frequency in the presynaptic neuron. We also investigate how noise in AP timing varies with different parameters, such as the probability of releases, the number of docking sites, the voltage threshold for AP firing, and the timescale of voltage decay. In summary, our results provide a systematic understanding of how stochastic mechanisms in neurotransmission enhance or impinge the precision of AP fringing times.

## I. INTRODUCTION

In the nervous system, communication between two neurons often occurs via a chemical synapse where action potential (AP)-triggered neurotransmitter release from the presynaptic neuron triggers an AP in the postsynaptic neuron. The efficacy for synaptic connections has been studied in several works [1]–[7] since cognitive processes such as learning, and memory depend on the synaptic efficacy. The concept of synaptic efficacy has an intuitive definition – the maximum amount of influence on a postsynaptic neuron from the presynaptic neuron(s). Previous works have shown that under some conditions, noise enhances the synaptic efficacy [8].

Several works investigated stochastic models for synapses at molecular and network levels, [9]–[11]. Among various models, the Leaky Integrate and Fire (LIF) models [12], [13] are commonly used. In LIF models, neurotransmitters released from the presynaptic neuron(s) alter the membrane potential of the postsynaptic neuron, and “leaky” refers to the fact that the potential can decay back to its resting state in the absence of synaptic inputs.

Here we apply the formulas of Stochastic Hybrid Systems (SHS) that effectively combine discrete and continuous random processes [14]–[28] to investigate how stochasticity in neurotransmitter release impacts the timing of postsynaptic AP generation via the LIF model. This work builds on our previous work that modeled the presynaptic vesicle turnover [29]-[30] to understand the downstream impact of vesicles dynamics on postsynaptic processes. More specially, we precisely quantify the stochastic dynamic of neuron’s membrane potential ***v***(*t*) that increases in jumps based on neurotransmitter-release events in the presynaptic neuron and decreases continuously in between events. Using the framework of first-passage times, where an AP is triggered in the postsynaptic neuron when potential crosses a threshold [9]–[11], we develop novel formulas quantifying both the mean and noise in the timing of AP firing.

Note that the synaptic connections in some neurons do not depend on the arrival of APs, and they signal through graded transmission [31]–[33]. In some other neurons, both mechanisms exist [34]–[36]. Here, we focus solely on the neurons that require APs to evoke neurotransmitter release. Applying the stochastic hybrid model [29]-[30] and LIF model [37], we first write the moment dynamics for neurotransmitter-filled vesicles ***n***(*t*) in the presynaptic neuron and the membrane potential ***v***(*t*) in the postsynaptic neuron at time *t*. Later on, these dynamics are solved exactly and used to determine the statistics of postsynaptic AP firing times. Our results quantify saturation levels in postsynaptic AP frequency as a function of model parameters and determine the limits of noise suppression in AP timing.

## II. Model formulation

The overall model connecting the release of vesicles from the presynaptic neuron to AP-triggering in the postsynaptic neuron is illustrated in Fig. 1. We start by first describing the presynaptic dynamics.

**Fig. 1.**
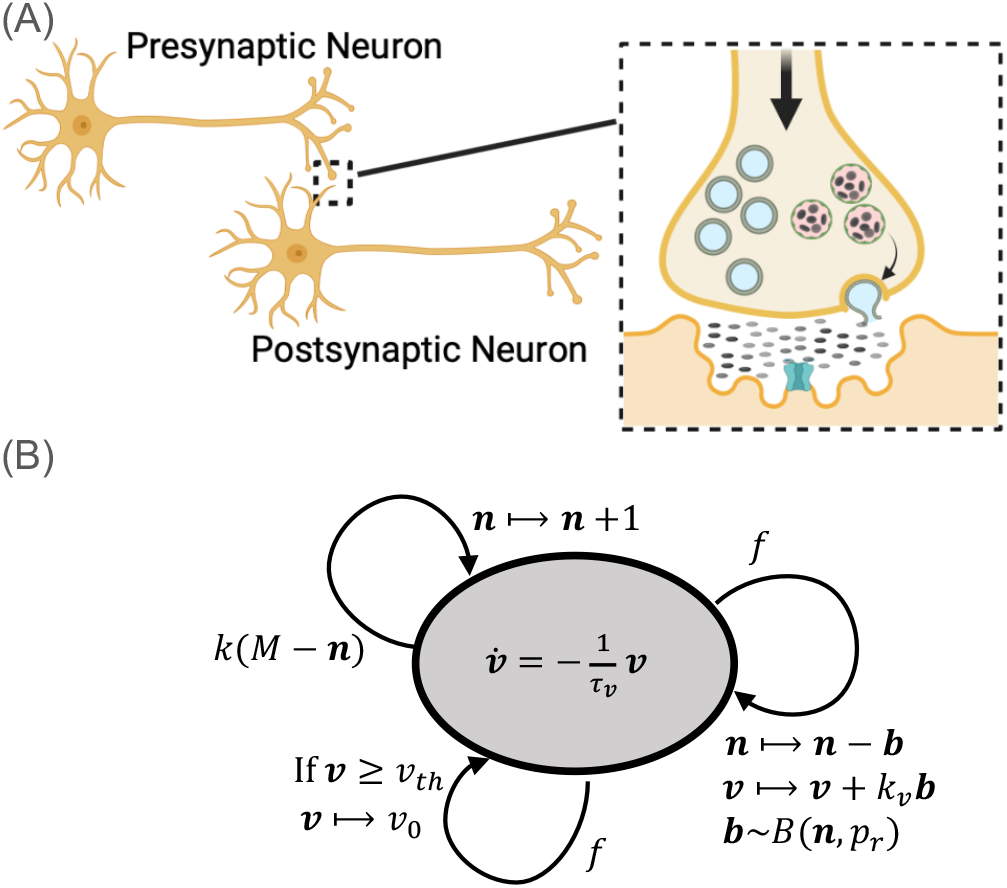
Schematic of two neurons communicating through a synapse and the corresponding SHS model. **(A)** In response to an AP reaching the axon terminal, docked vesicles are released emptying their neurotransmitter content into the synaptic cleft. Once a docked vesicle is released, the site becomes empty and each empty site gets refilled with vesicles. The neurotransmitters in the cleft regulate the opening of ion channels on the postsynaptic neuron’s membrane, resulting in the flow of charged ions (current) altering the membrane potential. **(B)** The SHS model of neurotransmission where the continuous dynamics within the circle represents the membrane potential ***v*** exponentially decaying over time [13]. The model has three different resets: The first reset corresponds to the arrival of AP in the presynaptic neuron with a rate *f* that causes ***b*** of the ***n*** docked vesicle to release. We assume ***b*** to follow a Binomial distribution with release probability *p*_*r*_. This event also corresponds to a jump in the membrane potential of the postsynaptic neuron by *k*_*v*_ ***b***. The second reset is the replenishment of docked vesicles in the presynaptic axon terminal with a rate *k*(*M* − ***n***). Finally, the third reset that is triggered when ***v*** ≥ *v*_*th*_ corresponds to an AP firing in the postsynaptic neuron that resets the membrane potential back to the resting potential *v*_0_.

### A. Presynaptic neuron

We assume that APs arrive at the axon terminal of the presynaptic neuron as per a Poisson process with a rate *f*. This corresponds to AP inter-arrival times being independent and identically distributed random variables following an exponential distribution with mean 1*/f*. Let the random process ***n***(*t*) denote the number of release-ready docked vesicles at time *t*. Upon AP arrival, ***b*** out of ***n*** vesicles are released

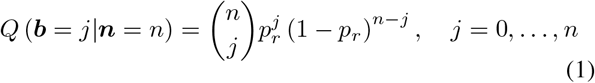

with ***b*** following a Binomial distribution, where *p*_*r*_ is release probability per vesicle. Since the Poisson arrival of an AP in the next infinitesimal time interval (*t, t* + *dt*) is *fdt*, the decrease in ***n*** by *j* ∈ {0, …, ***n***} vesicles is captured by the probabilistic event

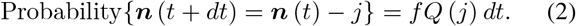

Between two successive APs, the number ***n*** builds up as a result of vesicle replenishment that occurs with rate *k* (*M* − ***n***(*t*)), where *M* is the number of docking sites (i.e., the maximum capacity for docked vesicles), and *k* is the refilling rate per site. This stochastic refilling is described by the probabilistic event

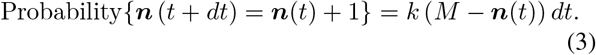

In summary, the continuous accumulation of ***n***(*t*) over time, and its depletion from binomial release occurring at discrete AP times are represented by events (3) and (2), respectively.

### B. Postsynaptic neuron

Assuming a rapid turnover of neurotransmitters in the synaptic cleft, a release of *j* vesicles leads to an instantaneous increase in the postsynaptic neuron’s membrane potential ***v***(*t*) by *k*_*v*_*j*, where *k*_*v*_ is a positive proportionality constant. As this increase is coupled with the arrival of APs in the presynaptic neuron, it can be directly combined with (2) to yield

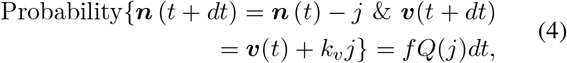

where *f Q*(*j*)*dt* is the probability of AP occurrence resulting in *j* released vesicles in the time interval (*t, t* + *dt*]. In between these discrete voltage jumps, ***v***(*t*) is assumed to decay via first-order kinetics

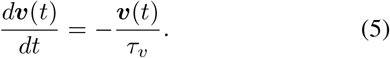

with *τ*_*v*_ quantifying the timescale of decay. Stimulation of the presynaptic neuron with a given frequency *f*, results in the buildup of membrane potential over time, and typical time traces of these random processes are shown in Fig. 2. An AP in the postsynaptic neuron is triggered when the membrane potential reaches a prescribed threshold *v*_*th*_. The timing of these postsynaptic APs can be mathematically formulated as the first passage time (FPT)

**Fig. 2.**
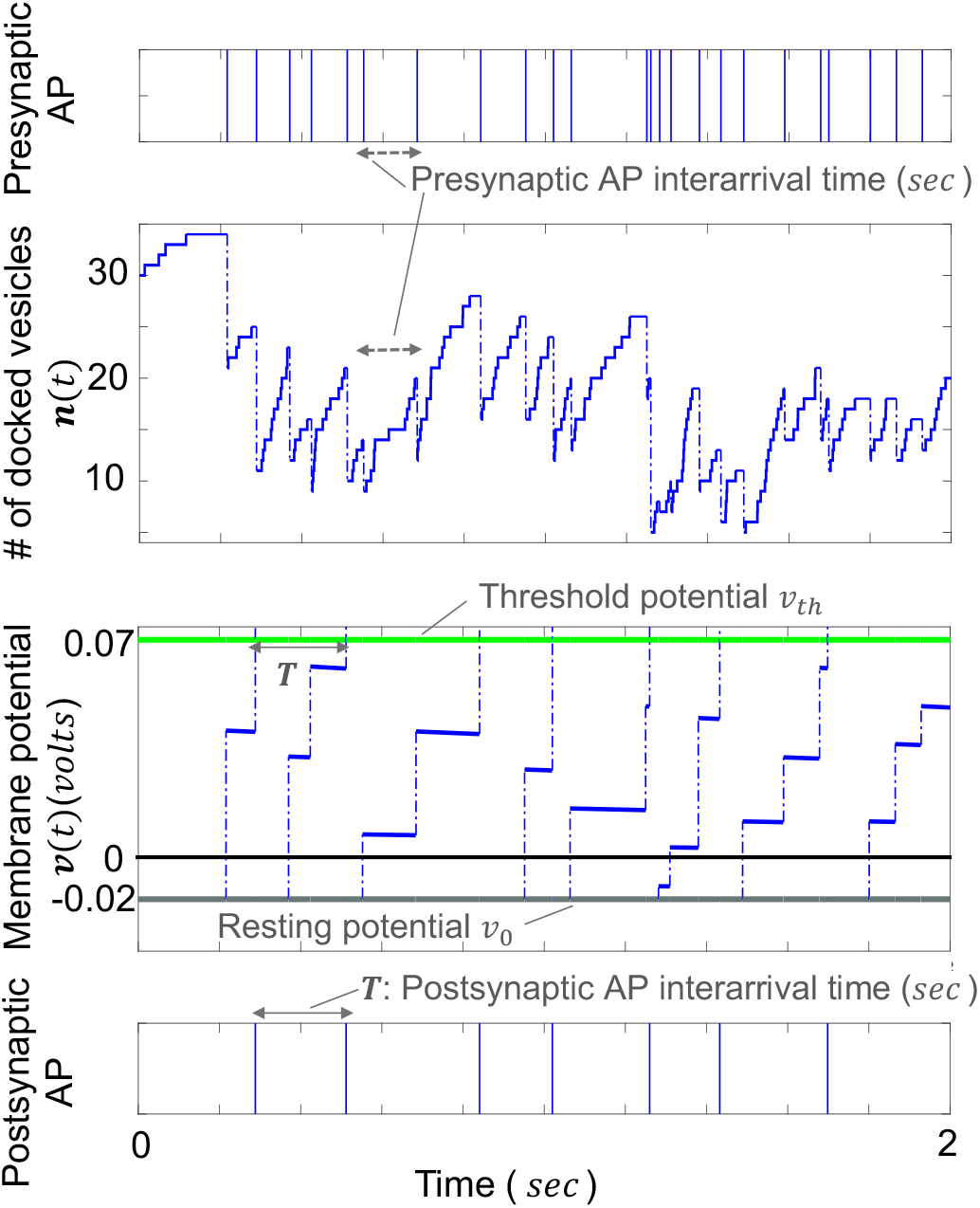
Sample trajectories for the number of docked vesicles *n*(*t*) in the presynaptic neuron and the postsynaptic neuron membrane potential *v*(*t*). The arrival of APs in the presynaptic neuron as per a Poisson process is shown on top together with the corresponding sample runs for ***n***(*t*) and ***v***(*t*) below. The membrane potential ***v***(*t*) increases by random jumps in response to presynaptic APs and decays continuously as per (5) in between APs. When ***v***(*t*) crosses the threshold level *v*_*th*_, an AP in the postsynaptic neuron is generated as shown in the bottom plot, and the membrane potential is reset to the resting value *v*_0_. For this plot, the model parameters are taken as *M* = 40, *k* = 5 *sec*^*−*1^, *f* = 10 *Hz, p*_*r*_ = 0.3, *τ*_*v*_ = 10 *sec* and *k*_*v*_ = 0.005 *volts*.

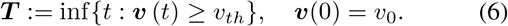

Here *v*_0_ denotes the resting membrane potential, which for convenience, we assume to be *v*_0_ = 0 volts. Once an AP is triggered in the postsynaptic neuron the membrane potential resets to *v*_0_. An essential goal of this investigation is to quantify the fluctuations in the first passage time ***T*** and determine how its mean ⟨***T***⟩ and noise vary as a function of different parameters. Throughout the paper, we use bold letters to indicate random variables/stochastic processes and, ⟨.⟩ and 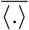 to denote the expected value and steady-state expected value respectively. For instance, the steady-state expected value of the stochastic process *x*(*t*) 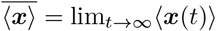 We also denote *f* and *F* = 1*/*⟨***T***⟩ the frequency of Aps in the presynaptic and postsynaptic neurons, respectively, and a complete list of model parameters are summarized in Table. 1.

**TABLE I.** Model parameters

| Parameter | Description |
| --- | --- |
| $v$ | Postsynaptic membrane potential (*volts*) |
| $n$ | Number of docked vesicles |
| $b$ | Number of released vesicles |
| $T$ | First passage time (FPT) (*sec*) |
| $f$ | Presynaptic AP frequency (*Hz*) |
| $F$ | Postsynaptic AP frequency (*Hz*) |
| $p_r$ | Release probability per site/vesicle |
| $k_v$ | Constant |
| $v_0$ | Membrane resting potential (*volts*) |
| $k$ | Refilling rate per vesicle (*sec*<sup>-1</sup>) |
| $M$ | Total number of vesicles |
| $v_{th}$ | Threshold potential (*volts*) |
| $\tau_v$ | Membrane time constant (*sec*) |

## III. The presynaptic neuron vesicles dynamics

The accuracy of the postsynaptic neuron’s response is critically dependent on stochasticity in neurotransmitters released from the presynaptic neuron [38]. In the previous section, we formulated a model that takes into account two important sources of such stochasticity arising from the replenishment and release of vesicles. Here we derive exact analytical expressions for the statistical moments of both the number of docked vesicle ***n***(*t*) in the presynaptic axon terminal and the number ***b*** released per AP.

Using standard tools from moment dynamics, the mean number of docked vesicles ⟨***n***(*t*)⟩ evolves as

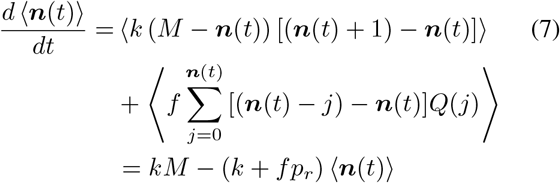

[39]–[42]. Given ⟨***n***(0)⟩ = *n*_0_, solving (7) yields

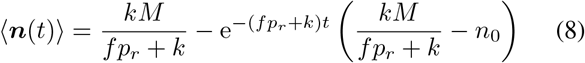

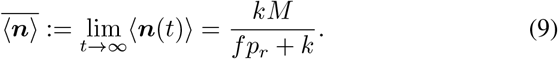

Assuming all docking sites are filled initially *n*_0_ = *M*, then presynaptic stimulation will cause a decrease in ⟨***n***(*t*)⟩ over time, and this vesicle depletion has often been referred to, in the literature, as synaptic depression [43], [44]. Fitting (8) to experimental data measuring the number of released vesicles over time upon presynaptic stimulation with a fixed frequency has been used to infer parameters across diverse synapses [45], [46]. Similar to (7), the differential equation describing the dynamics of the second-order moment is obtained as

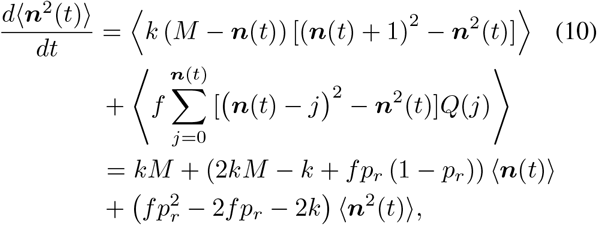

that results in the following steady-state solution

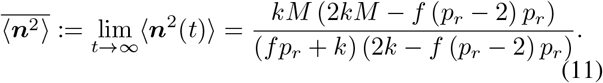

Using (9) and (11), we next quantify the extent of fluctuations in ***n***(*t*) by its squared coefficient of variation

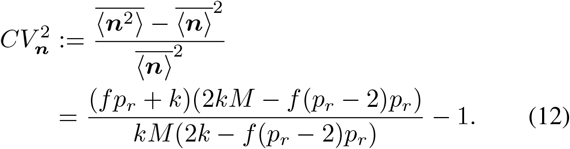

Since the number of released vesicles ***b*** conditioned on ***n*** follows a Binomial distribution as per (1), it is easy to see that at steady-state

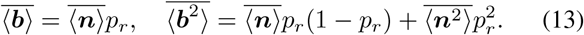

Using the above equation along with (9) and (11) yields the following coefficient of variation squared for ***b***

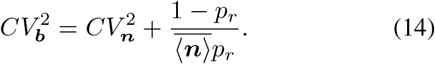

As expected, 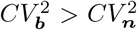 and 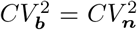 when *p*_*r*_ = 1 (i.e., all release-ready docked vesicles are released upon a presynaptic AP). Fig. 3 investigates the dependence of 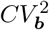 on key parameters: AP frequency *f*, release probability *p*_*r*_ and the vesicle refilling rate *k*. Intriguingly, our results show that 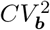 is minimized at an intermediate value of *p*_*r*_. In contrast, 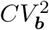 is a monotonically decreasing and increasing function of *k* and *f*, respectively (Fig. 3). How does randomness in the neurotransmitter-release process propagate downstream to impact AP triggering in the postsynaptic neuron?

**Fig. 3.**
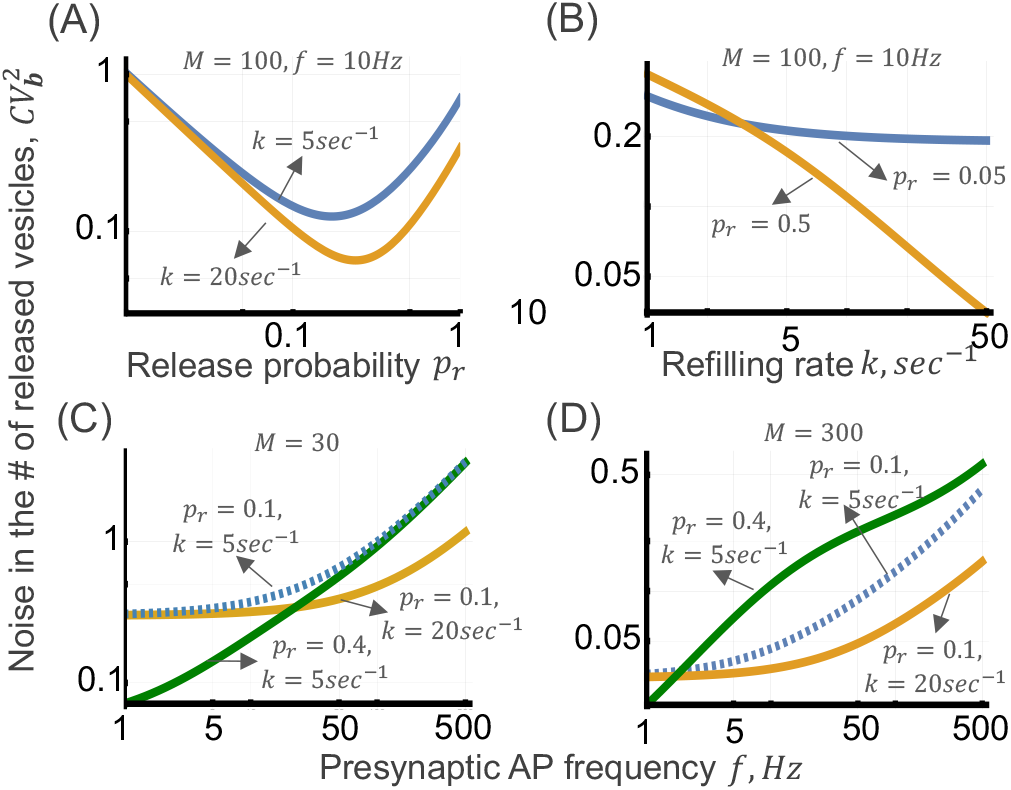
Stochasticity in the number of released vesicles per AP as a function of synaptic parameters. **(A)** The steady-state noise in the number of released vesicles 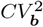 as given by (14) varies non-monotonically and is minimized at an intermediate value of *p*_*r*_. Parameters used to generate the lines are listed on the plot. **(B)** 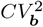 monotonically decreases with refilling rate *k*. Note that increasing *p*_*r*_ increases (decreases) 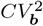 at low (high) values of *k*. **(C), (D)** 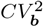 monotonically increases with AP frequency *f*. When *M* is small (left), increasing *p*_*r*_ from 0.1 to 0.4 for fixed *k* = 5 *sec*^*−*1^ attenuates 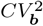. In contrast, when *M* is large (right), increasing *p*_*r*_ for fixed *k* = 5 *sec*^*−*1^ amplifies 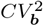.

## IV. Postsynaptic neuron AP formation

The released neurotransmitters result in an enhancement of the membrane potential, and potential buildup up to a threshold triggers an AP in the postsynaptic neuron. Recall from the model formulation that the time ***T*** between two postsynaptic APs is modeled as a first-passage-time (FPT) problem (6), and the goal here is to explore the output frequency *F* = 1*/*⟨***T***⟩ and stochasticity in ***T*** as measured by its coefficient of variation *CV*_***T***_. Our approach relies on first studying the impact of neurotransmitter release on the stochastic dynamics of the membrane potential ***v***(*t*), and then connecting fluctuations in the transient buildup of ***v***(*t*) to random fluctuations in ***T***.

### A. Stochastic dynamics of membrane potential

In the section, we derive formulas capturing the mean and noise in ***v***(*t*) over time. Given that the release of *j* vesicles results in an instantaneous increase in ***v***(*t*) by *k*_*v*_*j*, starting from the resting membrane potential of ⟨***v***(0)⟩ = *v*_0_ = 0 the average potential dynamics follows

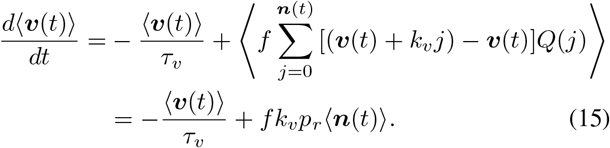

Assuming that the presynaptic dynamics is at equilibrium implying 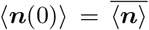 as per (9), the mean postsynaptic membrane potential increases exponentially followed by saturation as per first-order kinetics

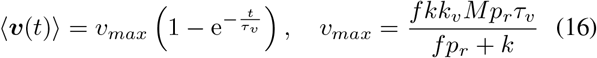

where *v*_*max*_ is the maximum level reached. Note that this maximum voltage is itself frequency-dependent and monotonically increases with *f* to reach

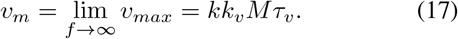

Since the threshold of AP firing *v*_*th*_ is generally much lower than *v*_*max*_, ***v***(*t*) never reaches close to *v*_*max*_ and is reset back to the resting potential once an AP is triggered. We next derive the dynamics for the second-order moments ⟨***n***(*t*)***v***(*t*)⟩ and ⟨***v***^2^(*t*)⟩ that are provided in the Appendix. Solving these linear dynamical systems with ⟨***n***(0)***v***(0)⟩ = 0, ⟨***v***^2^(0)⟩ = 0, and 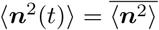 as given by (11), provides exact analytical formulas for the variance 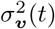 and the coefficient of variation squared 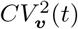 of the membrane potential over time. Due to space limitation, we only provide the formula for 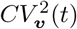 in (18) (see top of next page). In the limit of large input frequencies (*f* → ∞), 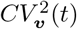 asymptotically follows

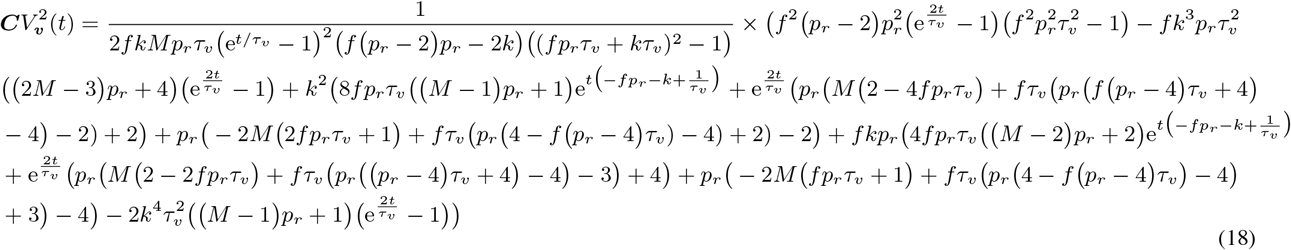

where

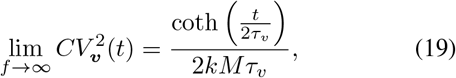

and (19) is invariant of the the release probability *p*_*r*_ and the constant *k*_*v*_. While the mean membrane potential increases over time, it turns out that 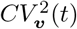 in (18) is a decreasing function of time. If we further take the limit *τ*_*v*_ → ∞ (i.e., there is no decay in membrane potential between successive vesicle-release events), then (19) simplifies to

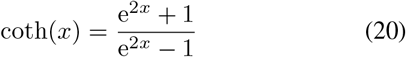

explicitly showing the inverse dependence with time.

### B. Postsynaptic neuron’s AP timing

Having derived an *exact* stochastic dynamics for the membrane potential, we now connect it to random fluctuations in ***T***. Assuming small fluctuations in ***v***(*t*) and *v*_*th*_ ≪ *v*_*max*_, the

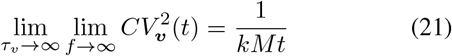

mean time ⟨***T***⟩ for AP firing is simply given by solving the mean potential reaching the threshold. Towards that end

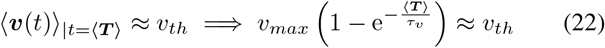

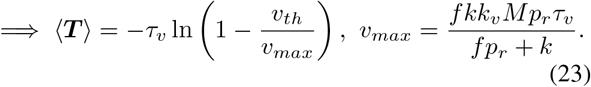

In this approximation regime, there exists a critical frequency *f*_*crit*_ that is obtained from solving *v*_*th*_ = *v*_*max*_, such that (23) is only defined for *f < f*_*crit*_ and ⟨***T***⟩ becomes unbounded as *f* → *f*_*crit*_. However, in the actual stochastic system low values of *f* will lead to large ⟨***T***⟩ implying low values of *F* = 1*/*(⟨***T***⟩. Moreover, in the limit of high frequency

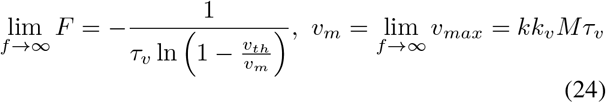

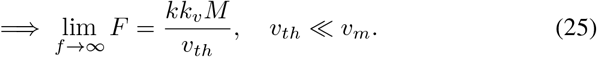

These results show that *F* is inversely proportional to the threshold *v*_*th*_, and is invariant of parameters such as *τ*_*v*_ and *p*_*r*_. We plot the postsynaptic AP frequency *F* versus the input frequency *f* in Fig. 4A, and the above analytical formula provides good agreement with frequencies obtained from stochastic simulation of the SHS model in Fig. 1.

**Fig. 4.**
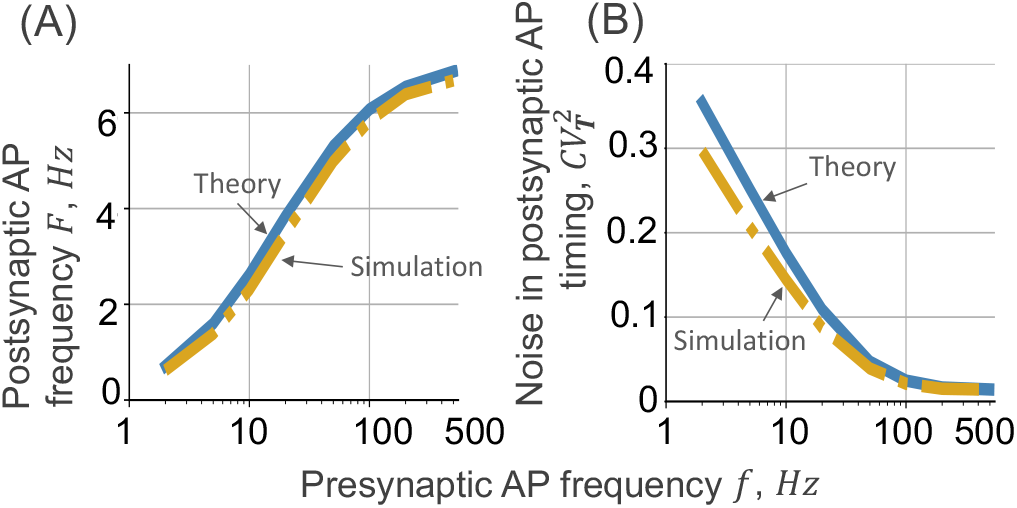
Noise in the postsynaptic AP timing decreases with increasing presynaptic AP frequency. **(A)** The output frequency *F* in the postsynaptic neuron as given by (23) is plotted as a function of input frequency *f*. The input-output plot follows an increasing sigmoidal that saturates at (25). **(B)** The noise 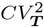 as derived in (28) as a function of *f*. Both *F* and 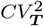 as predicted by the approximate formulas show good agreement with exact values obtained from stochastic simulations. Other parameters are taken as *M* = 100, *k* = 5 *sec*^*−*1^, *p*_*r*_ = 0.3, *v*_0_ = 0 *volts, v*_*th*_ = 0.07 *volts, k*_*v*_ = 0.001 *volts* and *τ*_*v*_ = 10 *sec* is chosen to be large so that the voltage decay between two APs is relatively small [47].

Next, we focus on quantifying the fluctuations in ***T***. Towards that end, we employ a useful geometric approximation

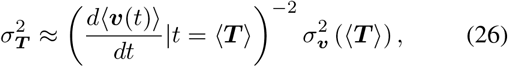

where the variance in ***T*** is connected to the variance in the membrane potential at time *t* = ⟨***T***⟩. The latter variance is further divided by the square of the slope of the mean potential buildup at time *t* = ⟨***T***⟩ implying a “flatter” approach to the threshold will lead to more fluctuations in the threshold-hitting time. This approximation has been widely used for studying timing in stochastic bio-molecular systems [48]–[52] and we use it here in the context of neurotransmission. This approximation yields the following formula for the squared coefficient of variation of ***T***

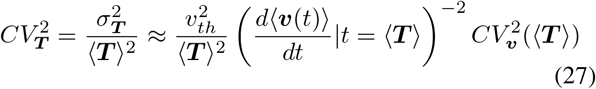

in which using (16) results in

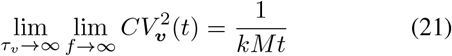

where 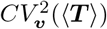 is given by (18) at *t* = ⟨***T***⟩ as in (22). This result highlights the points:

- 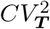 is proportional to the membrane potential fluctuation 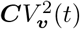 at time *t* = ⟨***T***⟩. Moreover, in the limit *v*_*th*_*/v*_*max*_ → 0, 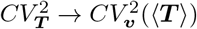.
- Consistent with stochastic simulation, 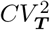 decreases with increasing frequency (Fig. 4B) and this is qualitatively different from behavior seen in Fig. 3, where 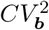 increases with *f*.

In the limit *f* → ∞, we substitute *F* from (24) in (28), where 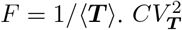 approaches

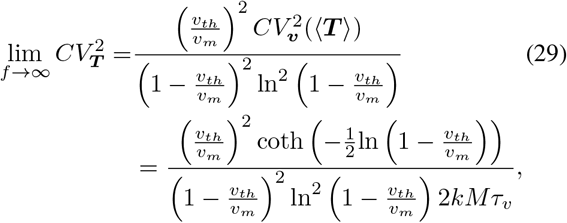

where *v*_*m*_ = lim_*f*→∞_ *v*_*max*_. Now further assuming *v*_*th* ≪_ *v*_*m*_, and using the fact that *x*^2^*/*(1 − *x*)^2^*/* ln^2^(1 − *x*) ≈ 1 + *x* for small *x*, (29) reduces to

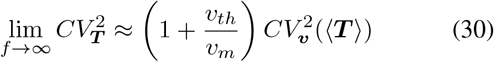

which after substituting (19) and (25) becomes

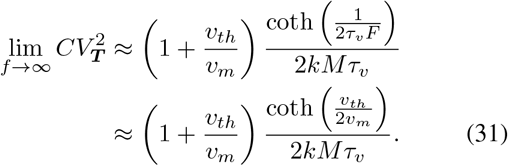

An interesting observation to note here is that 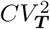 varies non-monotonically with *v*_*th*_*/v*_*m*_ (Fig. 5). In particular, 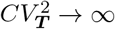 as *v*_*th*_*/v*_*m*_ → 0, and increasing the threshold first decreases 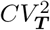 to reach a minimum, and then 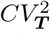 increases with increasing *v*_*th*_*/v*_*m*_. Note this increase is sharper in the original formula (29) where 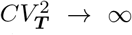 as *v*_*th*_ → *v*_*m*_ (Fig. 5). Also note from (31) that making the postsynaptic membrane potential more “leaky” by decreasing the value of *τ*_*v*_ enhances 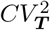.

**Fig. 5.**
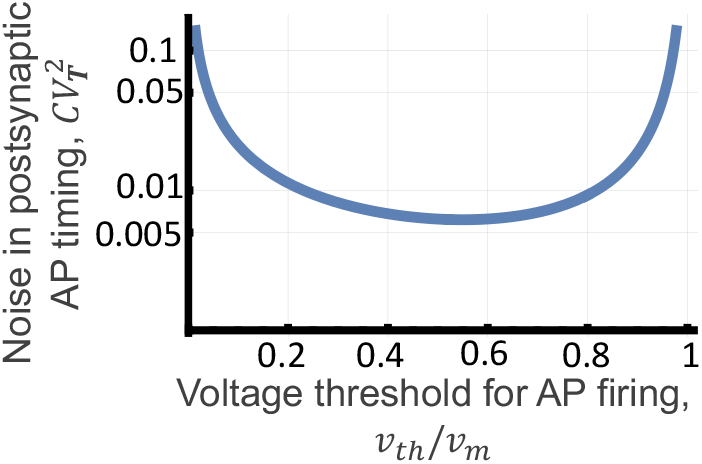
The noise in presynaptic AP timing 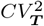 is minimized at an optimal threshold. Plot of 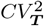 is given in (29) as a function of *v*_*th*_*/v*_*m*_ assuming 2*kMτ*_*v*_ = 10^3^.

### C. Frequency dependent release probability and refilling rate

So far, all the analyses have assumed constant values of the pre-synaptic parameters *k* and *p*_*r*_, which in reality itself depend on the frequency *f*. This mechanically occurs through the build of calcium in the axon terminal with increasing frequency that impacts vesicle refilling and release [53]–[55]. This effect can be modeled by simply having a frequency-dependent release probability *p*_*r*_(*f*) and refilling rate *k*(*f*) that follow Hill-type functions

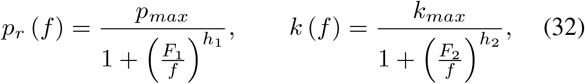

where *p*_*max*_ and *k*_*max*_ are the maximum values, *F*_1_ and *F*_2_ are numerically equal to the frequency at which *p*_*r*_ and *k* are half of their maximum values, respectively, and *h*_1_ and *h*_2_ are Hill coefficients. Fig. 6 illustrates how frequency-dependent parameters impact *F* and 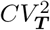 and essentially lead to curves that interpolate between the curves corresponding to fixed parameters.

**Fig. 6.**
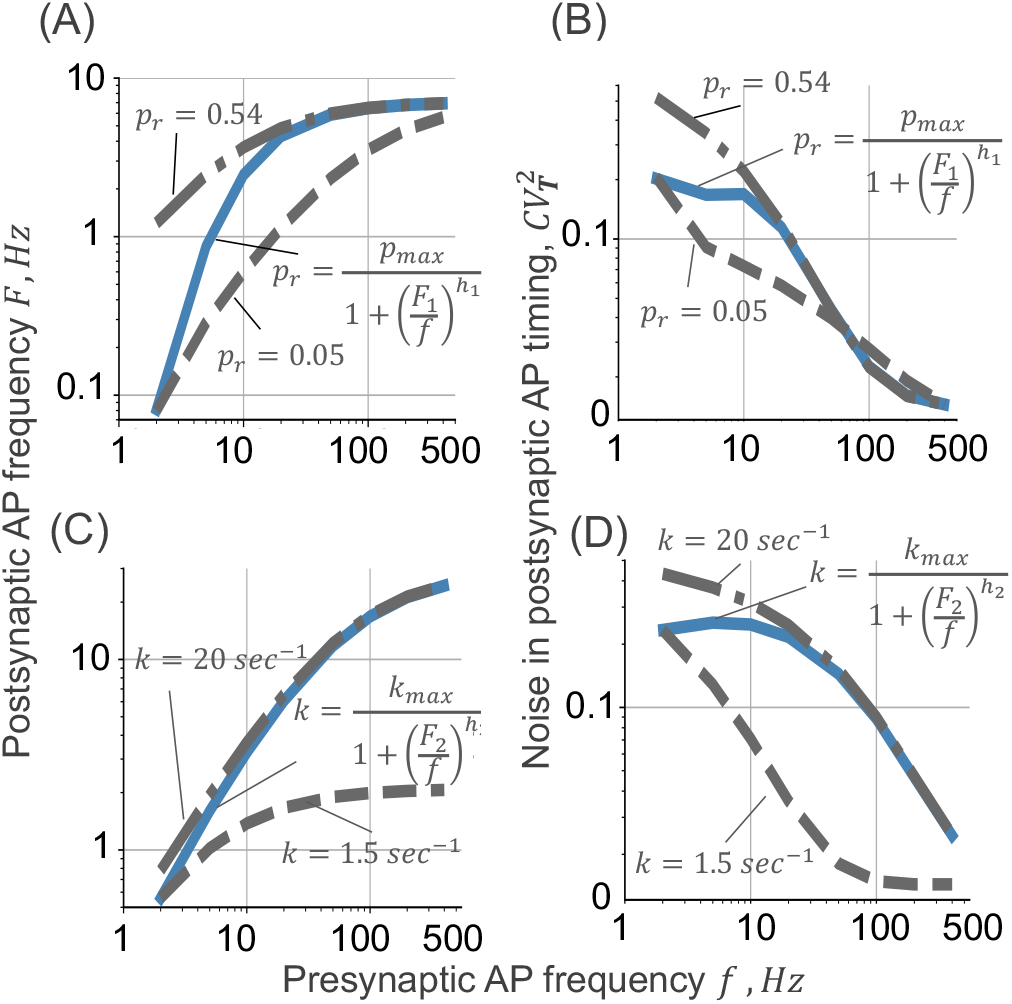
The postsynaptic AP frequency *F* and noise in AP timing 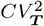 for frequency-dependent parameters. Plots of frequency *F* as given in (23) and noise (28) for fixed probability of release and vesicle refilling rates are compared with the case of frequency-dependent parameters as phenomenologically captured via (32). Parameters are chosen as *k* = 5 *sec*^*−*1^, *p*_*max*_ = 0.54, *F*_1_ = 10 *Hz, h*_1_ = 1.41, *k*_*max*_ = 20 *sec*^*−*1^, *F*_2_ = 10 *Hz, h*_2_ = 1.56 and *p*_*r*_ = 0.3, *M* = 100, *v*_0_ = 0 *volts, v*_*th*_ = 0.07 *volts, τ*_*v*_ = 10 *sec, k*_*v*_ = 0.001 *volts*.

## V. Conclusion

In this contribution, we have developed an SHS model to systematically analyze the interplay of presynaptic and postsynaptic stochastic processes. Our analysis developed closed-form expressions for the steady-state statistical moments of the number of docked vesicles ***n***, the number of vesicles released per AP ***b***, the postsynaptic membrane potential ***v*** and the postsynaptic AP timing ***T***. While the formulas for the first three random processes are exact, the formula for ***T*** is approximate as it involves the conversion of voltage-level fluctuations to threshold hitting-time fluctuations as per (26). On the presynaptic side, the results show noise in ***b*** to vary non-monotonically with the probability of release *p*_*r*_ (Fig. 4), and monotonically decrease with increasing presynaptic AP frequency *f*. This can be intuitively understood from the fact that in the limit *f* → ∞,

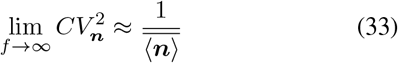

and similarly, it can be shown that

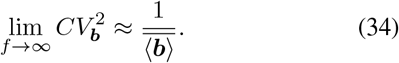

Thus, at high-frequency stimulation, both ***n*** and ***b*** follow Poisson statistics, and the coefficient of variation increases unboundedly as 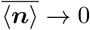 and 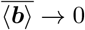 when *f* → ∞.

On the postsynaptic side, the output frequency saturation as *f* → ∞ is captured by (25) and this saturation is independent of *p*_*r*_. This point is exemplified in Fig. 6A, where the saturation levels are the same irrespective of fixed values of *p*_*r*_ and frequency-dependent *p*_*r*_. However, the maximum output frequency critically depends on the presynaptic vesicle refilling rate *k* (Fig. 6C). In some of our recent collaborations, we have uncovered that synapses involved in the auditory system have much higher values of *k* compared to other synapses [45], [46], and this may be important to enhance the dynamic range of operation needed for high-fidelity auditory functioning.

Our results also show that noise in AP generation timing decreases with increasing *f* (Figs. 4 & 6), and this is essentially a result of averaging multiple vesicle-release events that happen more effectively at higher values of *f*. This noise buffering is also evident from the fact that while the coefficient of variation of AP interarrival times is 1 in the presynaptic neuron due to the Poisson arrival assumption, the coefficient of variation of AP interarrival times on the postsynaptic side falls much below 1 (Figs. 4-6) The fundamental limits of noise suppression is given by (31) which is independent to *p*_*r*_, decreases with increasing *k*, and varies non-monotonically with the threshold *v*_*th*_. In the context of cell lysis from the random buildup of a toxin, there have been recent experiment validation of this U-shape dependence between noise in event timing and timing threshold [49]. It will be interesting to see if chemical synapses in-vivo also show similar behaviors and if they indeed tune the threshold to enhance precision in AP timing.

A part of our future work would be to compute the higherorder moments and the distribution for different random processes both at the presynaptic and postsynaptic sides. Another direction for our future work would be to expand the model to more complex models, including different types of vesicle pools, and consider multiple excitatory/inhibitory synaptic inputs on the postsynaptic neuron.

Appendix

The dynamics for (***n***(*t*)***v***(*t*)) and (***v***^2^(*t*)) follow

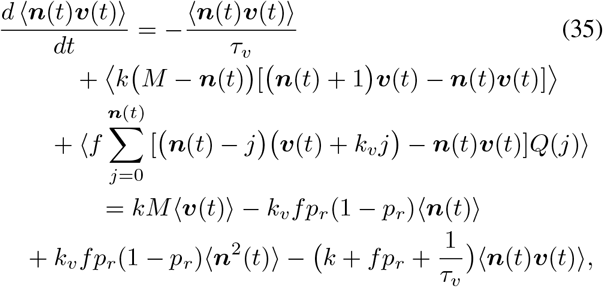

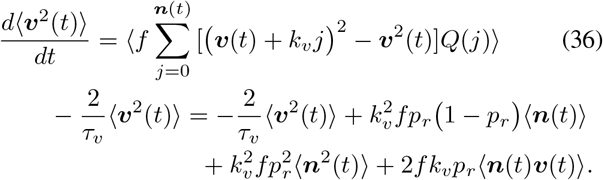

## ACKNOWLEDGMENT

This work is supported by NIH/NIDCD grant 1R01DC019268-01 and NSF grant ECCS-1711548.

